# Human Smc5/6 recognises transcription-generated positive DNA supercoils

**DOI:** 10.1101/2023.05.04.539344

**Authors:** Aurélie Diman, Gaël Panis, Cédric Castrogiovanni, Julien Prados, Bastien Baechler, Michel Strubin

## Abstract

Beyond its essential roles in ensuring faithful chromosome segregation and genomic stability, the human Smc5/6 complex acts as an antiviral factor. It binds to and impedes the transcription of extrachromosomal DNA templates; an ability which is lost upon chromosomal DNA integration. How the complex distinguishes among different DNA templates is unknown. Here we show that, in human cells, Smc5/6 preferentially binds to circular rather than linear extrachromosomal DNA. We further show that this binding is unlikely due to differences in the chromatin composition. Instead, the transcriptional process, per se, and more specifically the accumulation of DNA secondary structures known to be substrates for topoisomerases, is responsible for Smc5/6 recruitment. Those findings, in conjunction with our genome-wide Smc5/6 binding analysis showing that Smc5/6 localizes at few but highly transcribe chromosome loci, reveal a previously unforeseen role of Smc5/6 in DNA topology management during transcription.

## INTRODUCTION

The Smc5/6 complex belongs to the ring-shaped Structural Maintenance of Chromosomes (SMC) family complexes, which also includes the well-characterized cohesin (Smc1/3) and condensin (Smc2/4)^1^. These multi-subunit complexes, highly conserved in eukaryotes, are made of SMC heterodimers associated with a unique set of non-SMC-proteins (designated Nse1 to Nse4 in the case of Smc5/6). Powered by ATP, they entrap DNA molecules and contribute to chromosome architecture and dynamics^1, 2^. While the roles of cohesin and condensin in fundamental chromosomal transactions have been clearly established, the cellular functions specific to Smc5/6 remain to be clarified^1, 3^. The Smc5/6 complex has been linked to a wide range of cellular processes^4^ such as DNA replication^5^, DNA repair^6–9^, telomere maintenance^10^ and homologous recombination^11,12^. More recently, it has been identified as an antiviral factor targeting the hepatitis B virus (HBV)^13,14^. The binding of Smc5/6 to the circular HBV DNA genome impedes viral gene transcription and, thus, infection. To counteract the restriction activity of the complex, HBV expresses HBx, a protein that targets Smc5/6 for ubiquitin-mediated proteasomal degradation^13,14^. Smc5/6 restriction activity is not limited to HBV. Several studies have documented the ability of Smc5/6 to inhibit the transcription and/or replication of other human pathogenic viruses. These include human papillomavirus (HPV)^15,16^, herpes simplex (HSV-1)^17^, Kaposi’s sarcoma herpesvirus (KSHV)^18^, unintegrated human immunodeficiency virus (HIV-1)^19^, Epstein-Barr virus (EBV)^20^ and polyomavirus (SV40)^21^. The hallmarks of these viruses are the expression of inhibitory proteins antagonizing the Smc5/6 restriction activity and the maintenance of their genomes as a chromatinized extrachromosomal circular DNA within the nucleus of the infected cell^13,15–21^.

We previously reported that, as long as it remains extrachromosomal, the transcription of any reporter gene is silenced by Smc5/6. This occurs independently of the DNA sequence or the type of promoter used^13,22^. However, random chromosomal integration of the reporter gene safeguards it from Smc5/6-mediated restriction^13^. As shown by a recent structure–function analysis of the complex, extrachromosomal DNA (ecDNA) restriction is a property unique to Smc5/6, since neither cohesin nor condensin are involved^23^. The three-step restriction process involves the binding of Smc5/6 to ecDNA, their localization to Promyelocytic Leukemia Nuclear Bodies (PML-NBs) and subsequent transcriptional silencing through a yet unknown mechanism^23^. Since ecDNA entrapment by Smc5/6 is a prerequisite to its restriction activity, we investigated and characterized the DNA substrate requirements for Smc5/6 binding in human cells.

Using several extrachromosomal reporter gene constructs in combination with chromatin immunoprecipitation (ChIP) in non-transformed immortalized human retina pigment epithelial cells (hTERT-RPE1), we show that the chromatin composition resulting from different DNA uptake routes is not involved in ecDNA discrimination by Smc5/6. Instead, the circular nature of the ecDNA is essential for Smc5/6 recognition since a linear extrachromosomal construct escapes both Smc5/6 binding and restriction. We also found that the transcriptional process itself, and not the RNA polymerase II transcription machinery, is implicated in Smc5/6 recruitment onto ecDNA. Modulation of DNA topoisomerase activity, the enzymes resolving DNA topological stress occurring both during DNA replication and/or transcription^24^, provided evidence that Smc5/6 binding depends on the accumulation of transcription-driven topological constraints. Moreover, genome-wide analysis confirmed that the chromosomal association of Smc5/6 is also conditioned by transcription. Collectively our data suggest a previously unsuspected role for Smc5/6 in the management of DNA superhelical stress generated during transcription.

## RESULTS

### Smc5/6 restricts both non-replicated and replicated extrachromosomal DNA

For chromosomal genes, chromatin assembly mainly occurs during S phase and is tightly couple to DNA replication. In contrast, the chromatinization of exogenous DNA introduced into cells by transfection or viral transduction is uncoupled from DNA replication and can take place at any stage during the cell cycle^25–27^. To rule out the potential implication of an endogenous DNA modification, such as nucleosome composition or DNA methylation, in the specific recognition of extrachromosomal DNA by the Smc5/6 complex, we established a stable human cell line carrying an excisable chromosomal reporter construct that is not expressed when integrated into the chromosome (Figure 1A). The construct was engineered such that upon Cre recombinase expression, using a self-excising lentiviral vector^28^, the split Gaussia luciferase (GLuc) gene (GLuc-Nter and GLuc-Cter) flanked by two *LoxP* sites, embedded within a splice donor and acceptor sequences from an artificial intron, will form a circular extrachromosomal DNA molecule. After transcription, the *LoxP*-containing intron is spliced out and a functional GLuc mRNA is formed. To monitor Cre-mediated excision, a green fluorescent reporter (GFP) gene was inserted at the 3’ end of the construct while the constitutive EF1α promoter lies at the 5’ end. Upon excision, this configuration brings the GFP gene under the control of the EF1α promoter. Visualization of GFP-positive cells therefore provided a rapid and simple assessment of the recombination events (Figure 1B). Excision efficiency, as well as extrachromosomal circle formation, were further quantified by real-time quantitative PCR (qPCR). Using three primer sets spanning the construct (Figure S1A), we showed that whereas the signal for the GFP coding region did not change upon Cre-mediated excision (primer pair #G), as expected, the junction amplified by the primer pair #L almost completely disappeared. This coincided with the appearance of a new DNA junction, amplified by the primer pair #E, consistent with the generation of recombination-dependent extrachromosomal circles (Figure S1B).

**Figure 1.**
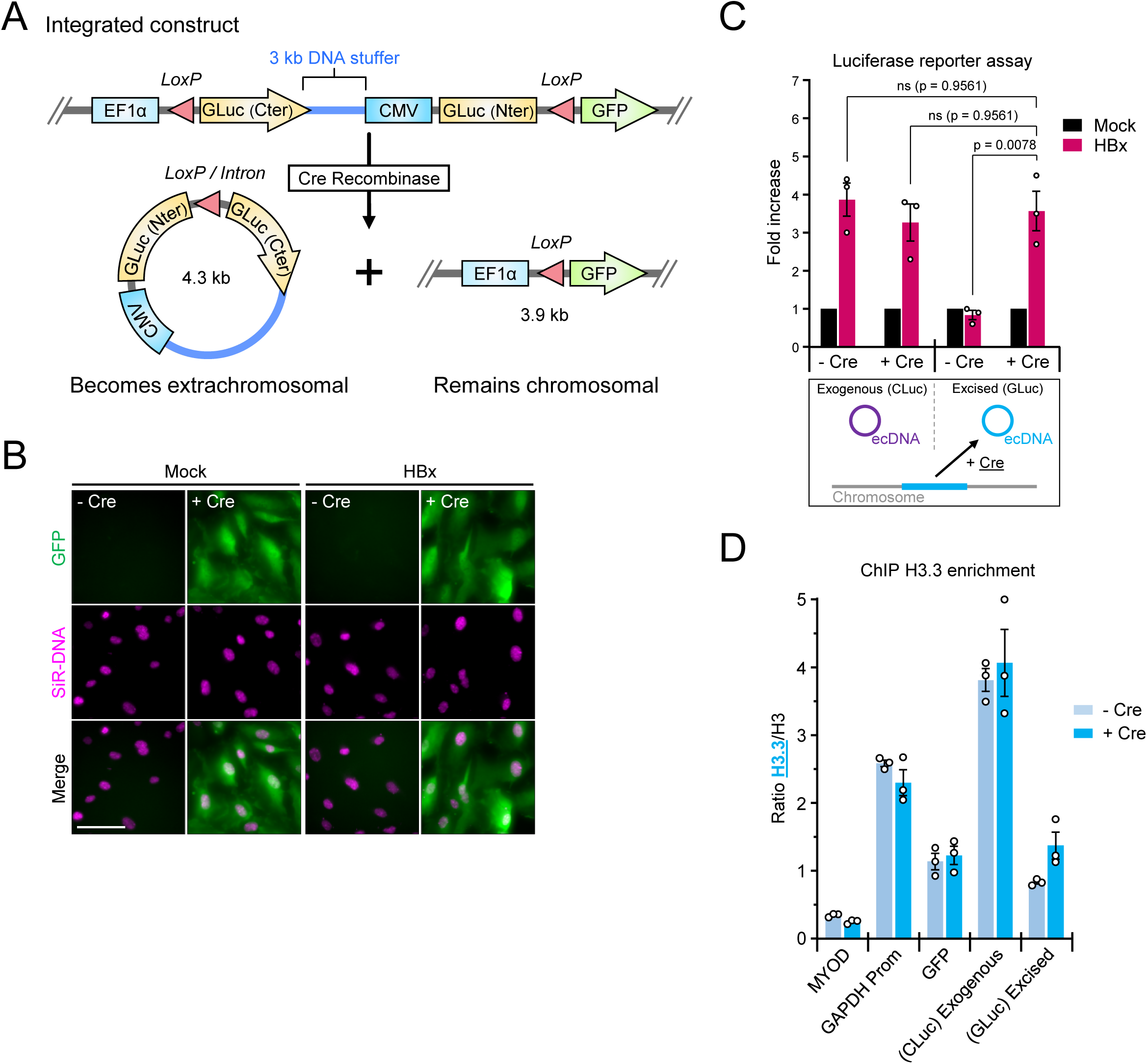
Non-replicated and replicated extrachromosomal DNA are both restricted by Smc5/6. (A) Schematic depiction of the genome integrated construct used to generate extrachromosomal [Gluc^circle^] and a chromosomally expressed GFP gene upon Cre/*loxP*-mediated excision. The yeast DNA stuffer is depicted in blue. (B) Live-cell representative images of Cre/*loxP*-mediated excision in hTERT-RPE1 cells containing the genomic excisable [Gluc^circle^] construct were co-transduced with lentiviruses containing either no gene insert (Mock) or HBx, plus or minus the Cre recombinase. Nuclei were stained with SiR-DNA. Scale bar, 100 μm. Data are representative of three independent experiments. (C) hTERT-RPE1 cells containing the genomic excisable [Gluc^circle^] construct (excised ecDNA) were co-transduced with lentiviruses containing either no gene insert (Mock) or HBx, plus or minus the Cre recombinase, together with an integrase-defective lentiviral Cypridina luciferase (CLuc) reporter construct (exogenous ecDNA). The luciferase assay was performed 2 days post-transduction. Luciferase activities are relative to their corresponding mock, which were set to 1. Data are means ± SEM of 3 independent experiments. Statistical analysis was performed using one-way ANOVA with Tukey’s multiple comparisons. (D) Same samples as in (C), histone binding at the indicated loci was monitored by ChIP using anti-H3 or anti-H3.3 antibodies. Values are expressed as a ratio of H3.3-enriched loci. Data are means ± SEM of 3 independent experiments. See also Figure S1

Measuring luciferase activity, we showed that GLuc gene expression from the excised extrachromosomal circle increased in a Cre-dependent fashion following Smc5/6 complex degradation by HBx. Transduction in the same cells of an extrachromosomal reporter construct of exogenous origin expressing a Cypridina luciferase gene (CLuc) delivered thanks to an integrase-defective lentivirus (IND64A) showed a similar increase in luciferase activity but independently of Cre expression as expected (Figure 1C). Similar results were obtained using two other cellular clones (Figure S1C).

Since both reporter constructs are responsive to the Smc5/6 complex to the same extent, we had to exclude that, despite their different origins, an identical histone content could explain their Smc5/6 behaviour. We therefore performed chromatin immunoprecipitation using anti-histone H3 and H3.3 antibodies followed by qPCR with specific primers. We confirmed the specificity of the anti-H3 and anti-H3.3 antibodies by comparing well-characterized genomic loci, the first exon of *MYOD* gene and *GAPDH* gene promoter, as positive controls for H3 or H3.3, respectively. We found that, compared to the excised extrachromosomal reporter, the exogenous reporter shows higher H3.3 occupancy, as expected for a chromatin assembled outside of the S phase (Figure 1D). Moreover, after Cre-mediated excision, the histone H3 and H3.3 occupancy does not change (Figure1D and S1D)^26,27^. This suggests that excision of the DNA circle has no major impact on its chromatin composition. Overall, these results demonstrate that Smc5/6 can restrict extrachromosomal reporter gene independently of their origin and chromatin composition.

### Smc5/6 preferentially binds and restricts circular DNA templates

The preferential entrapment of circular DNA molecules, *in vitro*, by the purified budding yeast Smc5/6 complex^29,30^, prompted us to ask whether a linear extrachromosomal DNA would be detected by the human Smc5/6 complex in a cellular context. For this, we modified an existing linear bacterial vector called pJazz®-OK^31^ and made it suitable for reporter gene expression in mammalian cells. The GLuc gene under the control of a cytomegalovirus promoter (CMV) was cloned into the vector (Figure 2A, upper panel). The covalently closed hairpin ends of the plasmid prevent its concatenation and its integration into the cellular chromosome. Taking advantage of two restriction enzyme sites (BssHII) present at both ends of the linear plasmid, we generated its circular counterpart thereby allowing a direct comparison of the two constructs that excluded any sequence bias (Figure 2A, lower panel). Luciferase assays following transient transfection with either the circular or linear reporter constructs, revealed that the circular DNA template was more strongly responsive to HBx compared to the linear template, which showed only weak stimulation (Figure 2B). Furthermore, and as expected, despite similar transfection efficiency as measured by qPCR analysis (Figure S2A), the basal expression levels of the linear construct approximated those measured for the circular DNA template in the presence of HBx (Figure S2B). We then examined whether the reduced stimulation of the linear reporter reflected a defect in Smc5/6 binding. ChIP experiments demonstrated, that indeed, Smc5/6 failed to stably associate with the linear extrachromosomal DNA (Figure 2C). Hence, linear extrachromosomal DNA escapes Smc5/6 entrapment and silencing.

**Figure 2.**
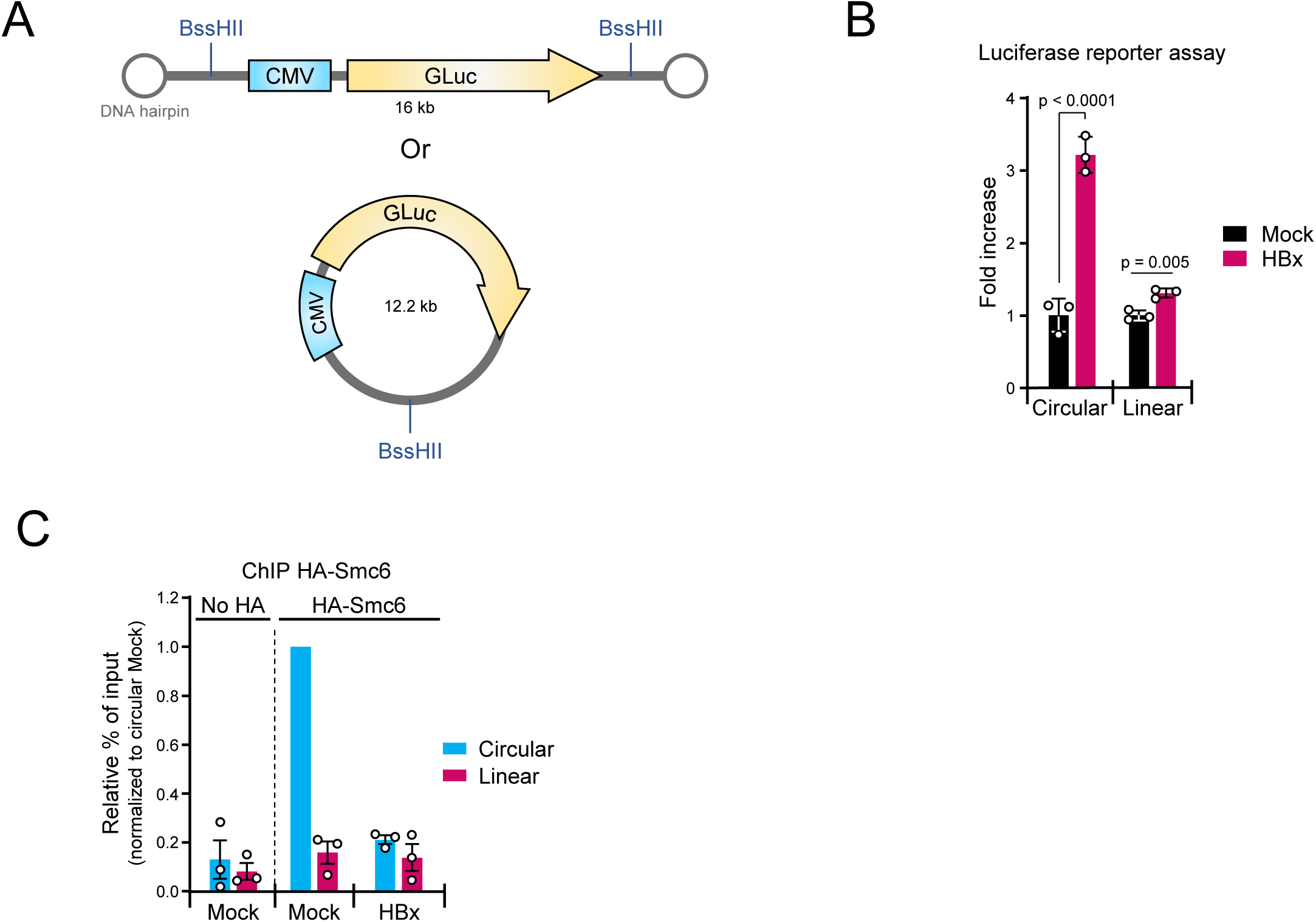
The binding and restriction activity of Smc5/6 are specific to circular DNA templates. (A) Schematic illustration of the linear pJAZZ-derived vector with its terminal DNA hairpin loops, containing a mammalian expression cassette. The CMV promoter drives the expression of the Gaussia Luciferase gene. The BssHII restriction sites used to convert the linear vector into its circular form are also indicated. (B) hTERT-RPE1 cells over-expressing the HA-tagged versions of Smc6 were transiently transfected with either a circular or a linear Gaussia luciferase reporter plasmid and then transduced with lentiviruses containing either no gene insert (Mock) or HBx. Luciferase activities were measured 2 days later and are indicated relative to their corresponding mock, which were set to 1. Data are means ± SEM of 3 independent experiments. Statistical analysis was performed using the Student’s t-test. (C) hTERT-RPE1 cells (No HA) or hTERT-RPE1 cells over-expressing HA-tagged version of Smc6 were transiently transfected with either a circular or a linear Gaussia luciferase reporter plasmid and then transduced with lentiviruses containing either no gene insert (Mock) or HBx. Anti-HA ChIP was performed 2 days later. The data are expressed as a percentage relative to the input normalized to the circular mock, which was set to 1. Data are means ± SEM of 3 independent experiments. See also Figure S2

### Smc5/6 binding to extrachromosomal DNA requires transcription but not RNA polymerase II

Chromosomal association of Smc5/6 has been reported to prevent accumulation of replication-induced DNA supercoiling^32^. Since the extrachromosomal reporter plasmids used in this study do not replicate, therefore excluding a role for replication in Smc5/6 binding, we investigate whether the topological stress induced by transcription promoted Smc5/6 loading onto extrachromosomal DNA. We performed time course experiments using two well-known transcription inhibitors. Cells were treated either with Actinomycin D (ACTD), which intercalates into the DNA helix thereby impeding the progression of the RNA polymerases^33^ or with Triptolide (TPT), which inhibits the transcription initiation step and induces proteasomal-dependent degradation of the RNA polymerase II (RNAP II)^34^. Both inhibitory treatments suppressed RNA synthesis, as confirmed by the reduced incorporation of the uridine analogue 5-ethynyluridine (EU), into newly transcribed RNA molecules (Figure S3)^35^. Using anti-Smc5 and anti-Nse4 antibodies, we further confirmed by western blot analysis that neither treatment alters the stability of the Smc5/6 complex, in contrast to what is observed upon HBx expression (Figure 3A and 3B). ChIP experiments using Smc6-HA-expressing hTERT RPE-1 cells, showed that transcriptional arrest resulted in a specific dissociation of Smc5/6 from extrachromosomal DNA while the level of histone H3 remained stable (Figure 3C and 3D).

**Figure 3.**
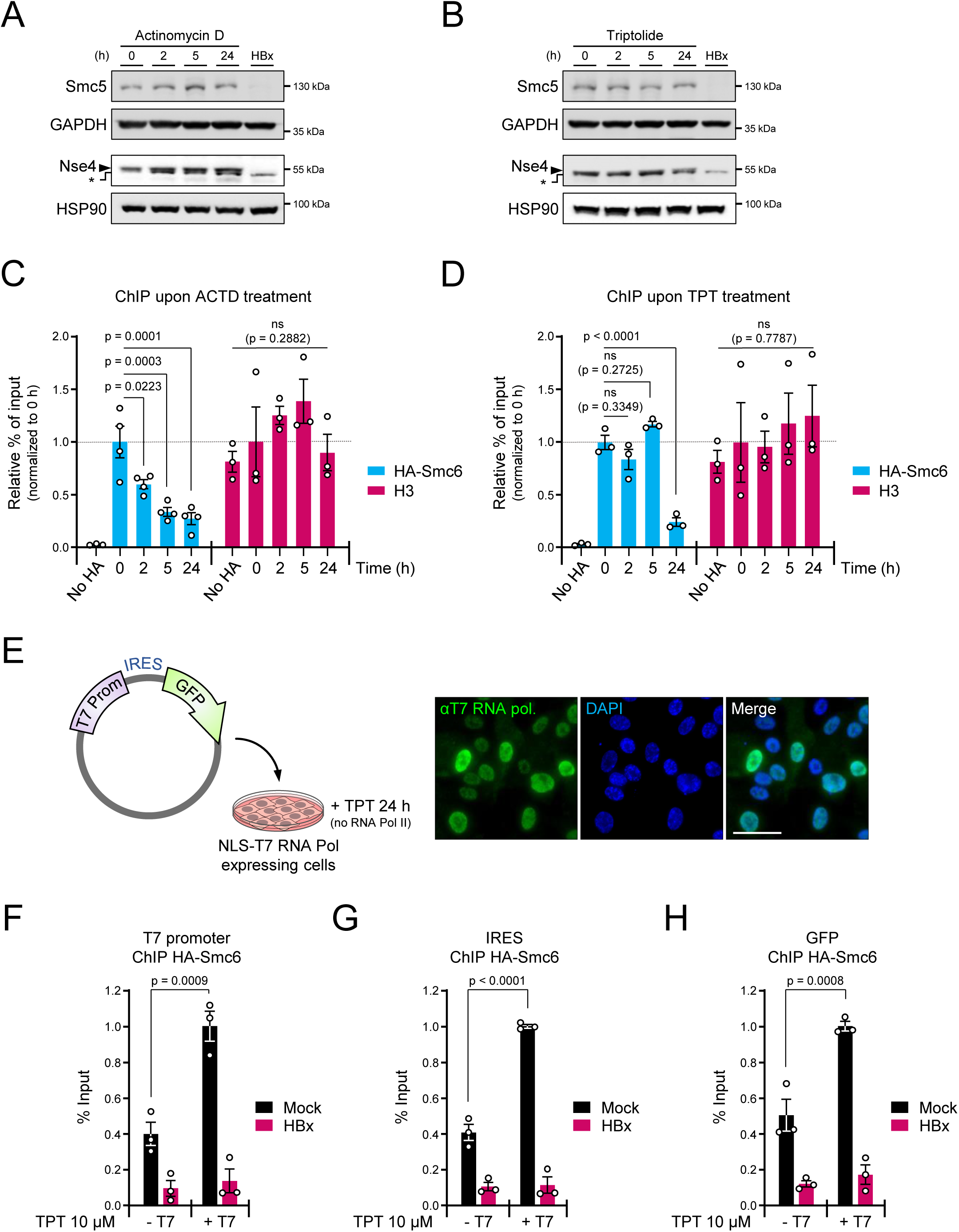
Recognition of extrachromosomal DNA by Smc5/6 is transcription-dependent but does not require RNA polymerase II. (A-B) Western blots showing Smc5 and Nse4 protein levels in protein extracts of hTERT-RPE1 cells over-expressing HA-Smc6 treated for the indicated times with (A) 10 μg/ml Actinomycin D (ACTD) or (B) 10 μM Triptolide (TPT). A protein extract of HBx treated cells was used as a control for Smc5/6 complex degradation. The asterisk (*) in the Nse4 blots indicates a non-specific band. Smc5 was used to assess the integrity of the Smc5/6 complex, because only a small fraction of the overexpressed HA-Smc6 is assembled into the Smc5/6 complex that binds DNA and is consequently degraded by HBx13,23. (C-D) hTERT-RPE1 cells (No HA) or hTERT-RPE1 cells over-expressing HA-tagged version of Smc6 were transduced with an integrase-defective lentiviral luciferase reporter construct and treated with (C) ACTD or (D) TPT for the indicated times before performing ChIP experiments with anti-HA (blues bars) or anti-H3 (pink bars). Data are expressed as a percentage relative to the input normalized to their corresponding 0 h time point, which was set to 1. Data are means ± SEM of 3 independent experiments. Statistical analysis was performed using one-way ANOVA with Tukey’s multiple comparisons. (E) Depiction of the experimental design (left panel). Immunofluorescence staining of hTERT-RPE1 HA-Smc6 cells transiently expressing the T7 RNA polymerase with a nuclear localization signal (NLS): confirmation of its nuclear localization (right panel). Nuclei were stained with DAPI. Scale bar, 50 μm. (F-H) HA-Smc6-expressing hTERT RPE-1 cells transiently expressing or not the T7 RNA pol were co-transduced with lentiviruses containing either no gene insert (Mock) or HBx, together with an integrase-defective lentiviral construct carrying a GFP gene under control of a T7 promoter and generating transcripts with an IRES sequence (Internal Ribosome Entry Site). At two days post transduction, cells were treated with 10 μM Triptolide (TPT) for 24h prior to the anti-HA ChIP experiments. Three different regions of the extrachromosomal construct, namely the T7 promoter (F), IRES (G) and GFP (H), were tested and compared to their respective mock values (minus T7 RNA pol). The data are expressed as a percentage relative to the input. Data are means ± SEM of 3 independent experiments. Statistical analysis was performed using one-way ANOVA with Tukey’s multiple comparisons. See also Figure S3

Since Smc5/6 association depends on transcription, we explored if the nature of the RNA polymerase II machinery was important for the recruitment of the complex to extrachromosomal DNA. An extrachromosomal construct carrying a GFP reporter gene under the control of the bacteriophage T7 RNA polymerase promoter was transduced in cells overexpressing a T7 RNA polymerase carrying a nuclear localisation signal (NLS-T7) (Figure 3E, left panel). The nuclear localization of the T7 RNA polymerase was confirmed by immunofluorescence microscopy (Figure 3E, right panel). Since it was previously reported that RNA pol II can drive transcription from a T7 promoter in mammalian cells^36^, cells were treated for 24h with TPT to block RNA Pol II transcription prior to ChIP experiments. We observed that T7 RNA polymerase driven transcription also leads to the recruitment of the Smc5/6 complex to extrachromosomal DNA (Figures 3F-3G). Altogether, those results highlight the notion that Smc5/6 binding is transcription-dependent but preclude that the recruitment occurs through interaction with the RNA pol II transcription machinery.

### Smc5/6 detects transcription-induced topological structures that are substrates for topoisomerases

Several studies have found that both the human and yeast Smc5/6 complexes, have a marked preference for DNA tertiary structures such as plectonemes that arise upon accumulation of DNA supercoils^30,37^. Our data indicating that circular DNA, as well as transcription, are required for Smc5/6 binding (Figures 2 and 3) prompted us to investigate the potential role of DNA topology in Smc5/6 recruitment to extrachromosomal DNA. We hypothesized that Smc5/6 dissociation upon transcriptional arrest was due to the activity of topoisomerases (Top) which function to dissipate transcription-generated supercoils^24^. Concomitant knock-downs of Top1, Top2A and Top2B was performed to prevent any functional compensation of Top2A by Top2B^38,39^. Western blot analysis failed to detect Top2A in hTERT RPE-1 cells under our experimental conditions (Figures 4A and 4C), but successful detection and knock-down validation was achieved in COLO320DM cells which exhibited higher Top2A expression levels (Figure S4). As expected, the simultaneous siRNA-mediated knock-down of Top1, Top2A and Top2B (Fig4A-S4) in combination with transcription inhibition, prevented the dissociation of Smc5/6 from the extrachromosomal DNA as shown by ChIP (Figure 4B). Furthermore, c-Myc overexpression, which has been shown to stimulate Top1 and Top2 relaxation activities through the formation of the “topoisome” complex^40^, led to Smc5/6 detachment, an event that could be counteracted by concomitant Top1, Top2A and Top2B siRNA-mediated depletion (Figures 4C-4D). Finally, to induce a targeted supercoil DNA relaxation of the ecDNA molecules, we overexpressed the myc-tagged DNA topoisomerase IB from Vaccinia virus fused to a NLS (Myc-NLS-vTopIB) (Figures 4E and 4F). This enzyme specifically recognizes the 5′-(C/T)CCTT-3′ DNA sequence^41,42^ present at three different locations over the ecDNA. ChIP experiments performed in these cells revealed a reduction in Smc5/6 ecDNA binding (Figure 4G) without an alteration in Smc5/6 complex stability (Figure 4F). Altogether, these results confirm the Smc5/6 DNA topology-dependent association.

**Figure 4.**
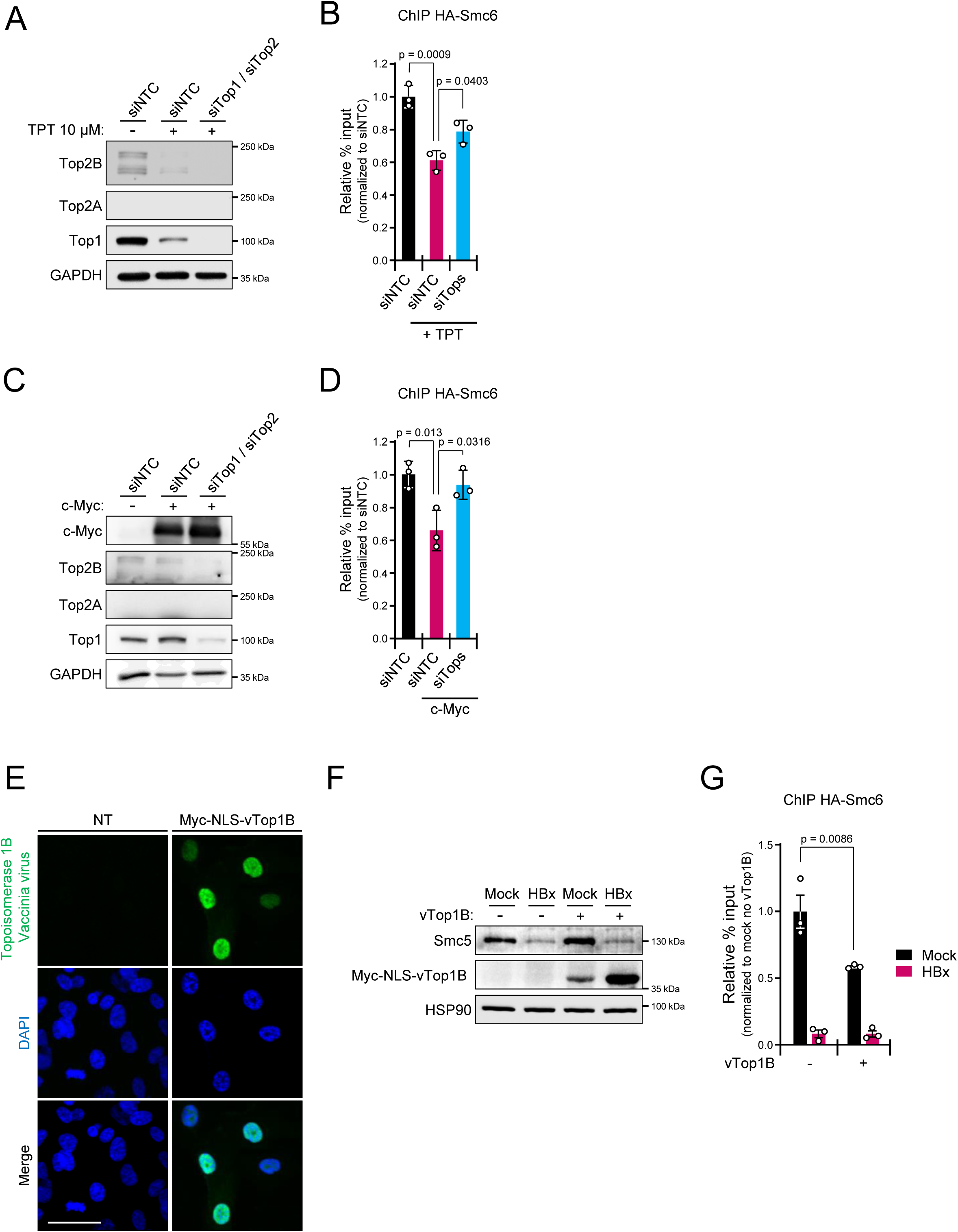
Smc5/6 recognizes topological structures arising during transcription that are substrates for topoisomerases. (A-B) HA-Smc6-expressing hTERT RPE-1 cells transduced with an integrase-defective lentiviral luciferase reporter construct were transfected with a non-targeting control siRNA (siNTC) or with siRNAs against topoisomerase 1 (siTop1) or topoisomerases 2A and 2B (siTop2) for 2 days before treatment or not with 10 μM Triptolide (TPT) for 24h prior to (A) Western blot analysis and (B) ChIP experiments using anti-HA. ChiP data are expressed as a percentage relative to the input normalized to the siNTC alone (or not treated with TPT), which was set to 1. Data are means ± SEM of 3 independent experiments. Statistical analysis was performed using one-way ANOVA with Tukey’s multiple comparisons. (C-D) HA-Smc6-expressing hTERT RPE-1 cells co-transduced with lentiviruses containing either no gene insert (Mock) or encoding the c-Myc gene, together with an integrase-defective lentiviral luciferase reporter construct were transfected with a non-targeting control siRNA (siNTC) or with siRNAs against topoisomerase 1 (siTop1) or topoisomerases 2A and 2B (siTop2). (C) Western blot and (D) ChIP using anti-HA were performed after 3 days. The ChiP data are expressed as a percentage relative to the input normalized to the siNTC alone, which was set to 1. Data are means ± SEM of 3 independent experiments. Statistical analysis was performed using one-way ANOVA with Tukey’s multiple comparisons. (E) Immunofluorescence staining of HA-Smc6-expressing hTERT RPE-1 cells transduced or not (NT) with lentiviruses encoding a N-terminal 3xMyc-tag-NLS topoisomerase 1B from vaccinia virus (Myc-NLS-vTop1B) confirming its nuclear localization. Nuclei were stained with DAPI. Scale bar, 50 μm. (F-G) HA-Smc6-expressing hTERT RPE-1 cells transiently expressing or not Myc-NLS-vTop1B, transduced with an integrase-defective lentiviral luciferase reporter, were co-transduced with lentiviruses containing either no gene insert (Mock) or HBx. (F) Western blots using an anti-Myc confirmed the expression of Myc-NLS-vTop1B with no impact on Smc5/6 complex integrity as demonstrated by Smc5 protein levels. (G) ChIP with anti-HA were performed after 3 days. ChiP data are expressed as a percentage relative to the input normalized to no Topo1V mock, which was set to 1. Data are means ± SEM of 3 independent experiments. Statistical analysis was performed using one-way ANOVA with Tukey’s multiple comparisons. See also Figure S4

### Smc5/6 associates with positive DNA supercoils in human cells

Having shown that the accumulation of topological constraints due to transcription is responsible for the recruitment of Smc5/6 to ecDNAs, we asked if our results could recapitulate the Smc5/6 binding pattern on a genomic scale. Although Smc5/6 ChIP sequencing (ChIP-seq) data are publicly available for *S.cerevisiae*^43,44^ *and Mus.musculus* ^45^, to our knowledge, there is no such data reported for the human complex. Therefore, to gain more insights into the genome-wide Smc5/6 DNA-binding profile, we generated ChIP-seq data from HA-Smc6 and compared them to untagged Smc6 control cells. Our ChIP-seq results confirmed the ChIP-qPCR results with a 63-fold enrichment of reads mapping on the extrachromosomal GLuc ORF in the HA-Smc6 samples (Figure S5A). In addition, we identified a total of 41 binding sites for Smc5/6 throughout the human genome with a significant 2.5-fold enrichment compared to the no HA control cells (Figure 5A DMSO and No HA). To test if the chromosomal association of Smc5/6 was also transcription-dependent, we performed ChIP-seq experiments following TPT treatment. In agreement with our observations previously made on extrachromosomal DNA (Figure S5A), transcriptional arrest largely abrogated Smc5/6 binding on chromosomal DNA (Figure 5A TPT). Surprisingly, the Red Fluorescent Protein reporter gene (RFP) present in the HA-Smc6 construct integrated into the chromosome also displayed Smc5/6 transcription-dependent binding (Figure S5B) but is not restricted by the complex, as previously reported^13^. To investigate in more detail the chromosomal relationship existing between Smc5/6 and the transcription process, we also performed ChIP-seq for RPB1, the largest RNAP II subunit^46^, under the same experimental conditions. Heatmaps of the Smc5/6 bound-regions and RNAP II binding sites within 4 kb around the Smc5/6 peak summit (Figures 5A-5B) revealed a perfect colocalization between the Smc5/6 enriched-loci (78%) and the heavily bound RNAP II regions (Figure S5C). Comparison with publicly available RNA-seq profiles for RPE cells^47^, confirmed that those regions corresponded to hyperactive transcription sites (Figure S5D). Based on our previous observations, we decided to compare our Smc5/6-seq results with the published genome-wide mapping of Top1^48^ and Top2 activity^49^. We found that the Smc5/6 bound-regions perfectly overlapped with the Top1/Top2 bound loci along the transcription units, suggesting the presence of DNA supercoils at those loci (Figure 5C). Interestingly, careful analysis of the Smc5/6 bound-regions identified a statistically significant enrichment of Smc5/6 downstream of the highly transcribe genes as exemplified by the depicted genomic regions (Figure 5C). Altogether, these results point to a model where Smc5/6 preferentially binds positive DNA supercoils generated during transcription and accumulating at the 3’ends of highly transcribed genes.

**Figure 5.**
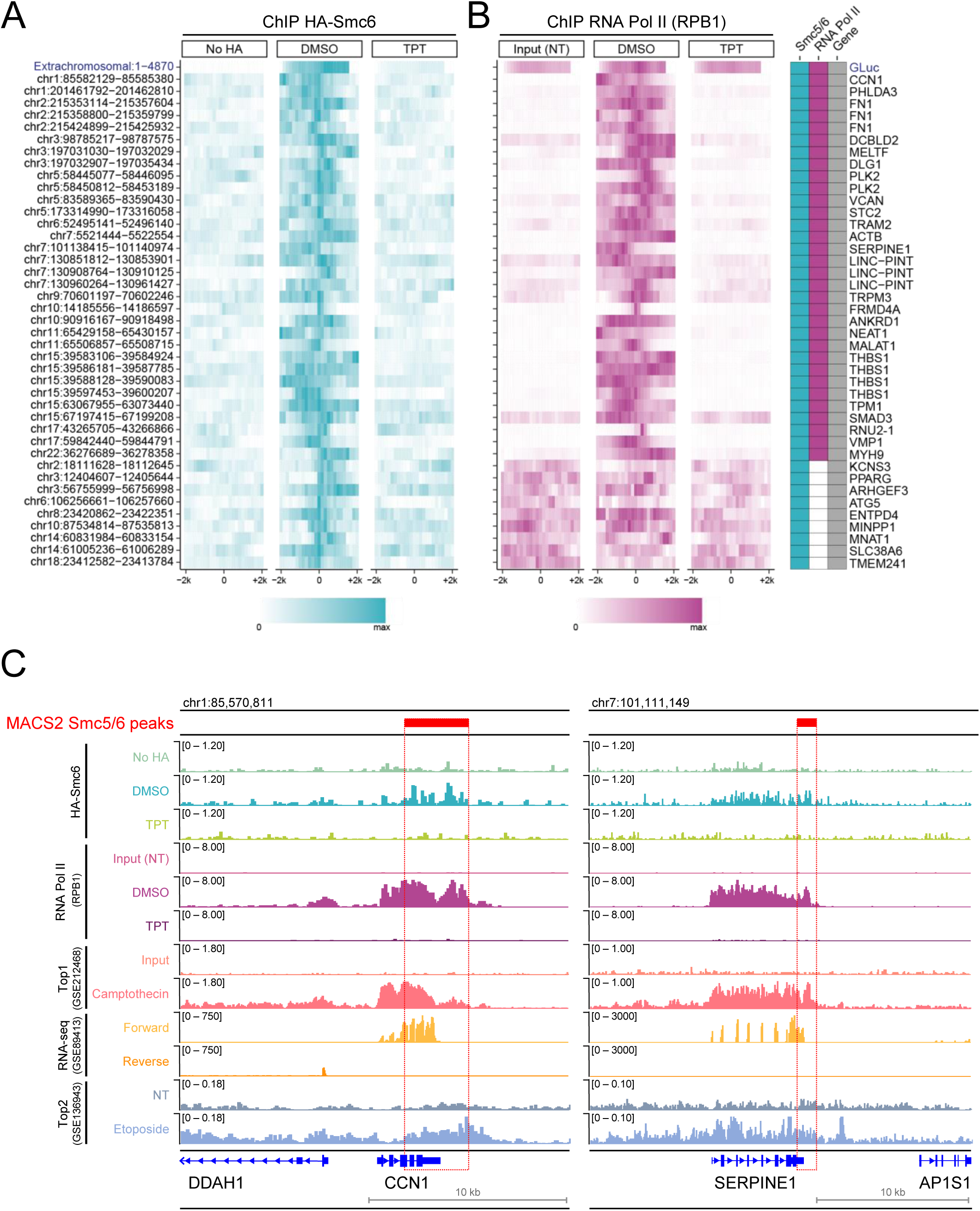
In human cells, Smc5/6 binds to positively supercoiled DNA. (A) Heatmaps of the ChIP-seq read depth around 41 identified Smc5/6 complex binding sites: in hTERT-RPE1 cells (No HA) (left panel); hTERT-RPE1 cells over-expressing a HA-tagged version of Smc6 treated either with DMSO (middle panel); and 10 μM Triptolide (TPT) (right panel) for 24h before HA-ChIP. Cells were transduced with an integrase-defective lentiviral luciferase reporter construct (GLuc). Rows represent Smc5/6 binding sites ±2 kb around the peak summit, ordered by chromosome number and according to the presence or absence of RNA Pol II as determined by RPB1-ChIP-seq (B). Only peaks statistically detected using MACS2 software analysis (2.5 fold enrichment, 0.05 q-value) are depicted. The colour scale represents the ChIP-seq normalized read depth (RPM) row-scaled identically across the 3 samples, with mapped reads virtually resized to 1 kb-length and looking at each genomic position for the amount of overlap between forward- and reverse-stranded reads. (B) Heatmaps of RNA pol II ChIP-seq peaks in hTERT-RPE1 cells over-expressing a HA-tagged version of Smc6 treated either with DMSO (DMSO), 10 μM Triptolide (TPT) or the corresponding input (NT), for 24h before RPB1-ChIP. Cells were transduced with an integrase-defective lentiviral luciferase reporter construct (GLuc). Rows represent RNA pol II binding sites ±2 kb around the Smc5/6 peak summit, ordered by chromosome number and according to the presence or absence of RNA Pol II as determined by the RPB1-ChIP-seq results. The colour scale represents the ChIP-seq normalized read depth (RPM) row-scaled identically across the 3 samples, with mapped reads virtually resized to 1 kb-length and looking at each genomic position for the amount of overlap between forward- and reverse-stranded reads. (C) Integrative Genomics Viewer (IGV) track screenshots from 2 representative genomic loci. The location of the Smc5/6 peaks identified by MACS2 are depicted with the red line. Tracks 1 to 3 (No Ha, DMSO, TPT) represent Smc5/6 ChIP-seq data for the corresponding samples and tracks 4 to 6 (Input NT, DMSO, TPT) represent RNA pol II ChIP-seq data for the corresponding samples. The subsequent tracks, Top1 (GSE212468)^48^, RNA-seq (GSE89413)^47^ and Top2 (GSE136943)^49^ depict publicly available data obtained for hTERT-RPE1 cells. Scales refer to the signal range in individual genome tracks. See also Figure S5

## DISCUSSION

The unexpected discovery that Smc5/6, besides its crucial role in maintaining genome stability, also acts as a transcriptional repressor that specifically targets extrachromosomal DNA, raises an important question^13,14,22^: How can a host genome architectural factor distinguish between chromosomal and extrachromosomal DNA inside the nucleus? As a first step towards elucidating the mechanism that makes Smc5/6-mediated transcriptional suppression specific to ecDNA, we characterized the DNA requirements for Smc5/6 binding. Using an excisable reporter construct, we show that the Smc5/6 complex does not discriminate ecDNA molecules based on their chromosomal or non-chromosomal origin. Since we showed that both, non-replicated and replicated extrachromosomal DNA, are restricted by Smc5/6, it precludes that specific chromatin marks, such as DNA methylation or histone composition, are involved in Smc5/6 specific DNA recognition. This is in agreement with recent findings showing that Smc5/6 has the ability to restrict EBV, HPV and KSHV whose viral genomes are maintained as extrachromosomal DNA packaged with chromatin in a manner similar to that of the host chromosome^15,16,18,20^. Instead, we have shown that Smc5/6 preferentially targets circular ecDNA molecules while linear ecDNA escapes Smc5/6 topological entrapment. These results are in agreement with *in vitro* pull-down experiments demonstrating the salt-stable binding of the budding yeast Smc5/6 complex to circular but not linear DNA templates^29,50^. We hypothesized that the helical topology of DNA could be a critical determinant for Smc5/6 loading. A linear DNA molecule can dissipate topological stress simply by spinning around its own helical axis, whereas a covalently closed circular DNA accumulates high levels of superhelical tension, building-up higher order DNA structures such as plectonemes^51^. The transcribing RNA polymerase is a potent generator of DNA supercoils^52^. According to the twin supercoiling domain model, the torque imposed on the DNA continuously generates negative supercoils behind and positive supercoils ahead of the RNA polymerase^53^. Our results showing that the transcription process per se, rather than components of the RNA pol II transcription machinery, is responsible for Smc5/6 recruitment onto circular ecDNA suggest an involvement of the DNA superhelicity. Supporting the DNA topology-dependent association of Smc5/6, we show that preventing supercoil removal by knocking down cellular topoisomerases counteracts the detachment of Smc5/6 observed after transcriptional switch-off. These findings were corroborated by the demonstration that Smc5/6 affinity for circular ecDNA was decreased upon reduction of their supercoil levels induced either by stimulation of endogenous topoisomerase activity or by overexpression of a viral topoisomerase.

Unexpectedly, our ChIP-seq analyses revealed a genome-wide interdependency between topological stress accumulation due to transcription and Smc5/6 binding at multiple, but nevertheless a restricted number of DNA loci. This suggests that the Smc5/6 complex is required in specific transcriptional scenarios because not all the transcribe loci were bound by the complex, but only those with elevated transcriptional levels. Therefore, we envision a model (Figure 6) in which the very high transcriptional output of certain genes leads to an abnormally high level of DNA supercoil accumulation, generating DNA secondary structures that ultimately result in the recruitment of the Smc5/6 complex. This model supports the recent findings of *in vitro* biophysical experiments made with the purified complex showing the preferential binding of Smc5/6 to structured DNA such as plectonemic substrates^30,37^. Although *in vitro* studies did not establish a clear preference of Smc5/6 for DNA supercoils based on their chirality, our results, suggest that *in vivo*, the Smc5/6 complex may preferentially bind to supercoiled DNA with positive handedness, as it is often found at 3’ ends of highly transcribed genes. Moreover, since Smc5/6 has been proposed to not only bind but also stabilize DNA plectonemes^30^, we advance a scenario in which Smc5/6 would trap and gather the excessive buildup of positive supercoils generated in front of the transcription machinery. The Smc5/6-mediated local plectoneme containment could either prevent supercoil propagation, which could inhibit subsequent transcription at neighboring genes, or promote the effective interaction between the DNA secondary structures and the topoisomerases or a combination of both. Collectively, our data suggest that by sensing and monitoring the levels of DNA torque *in vivo*, Smc5/6 would act as a topological insulator inhibiting the diffusion of transcription-induced positive supercoils. Future studies should be aimed at addressing *in vivo* the interplay between DNA supercoils, Smc5/6 and the topoisomerases.

**Figure 6.**
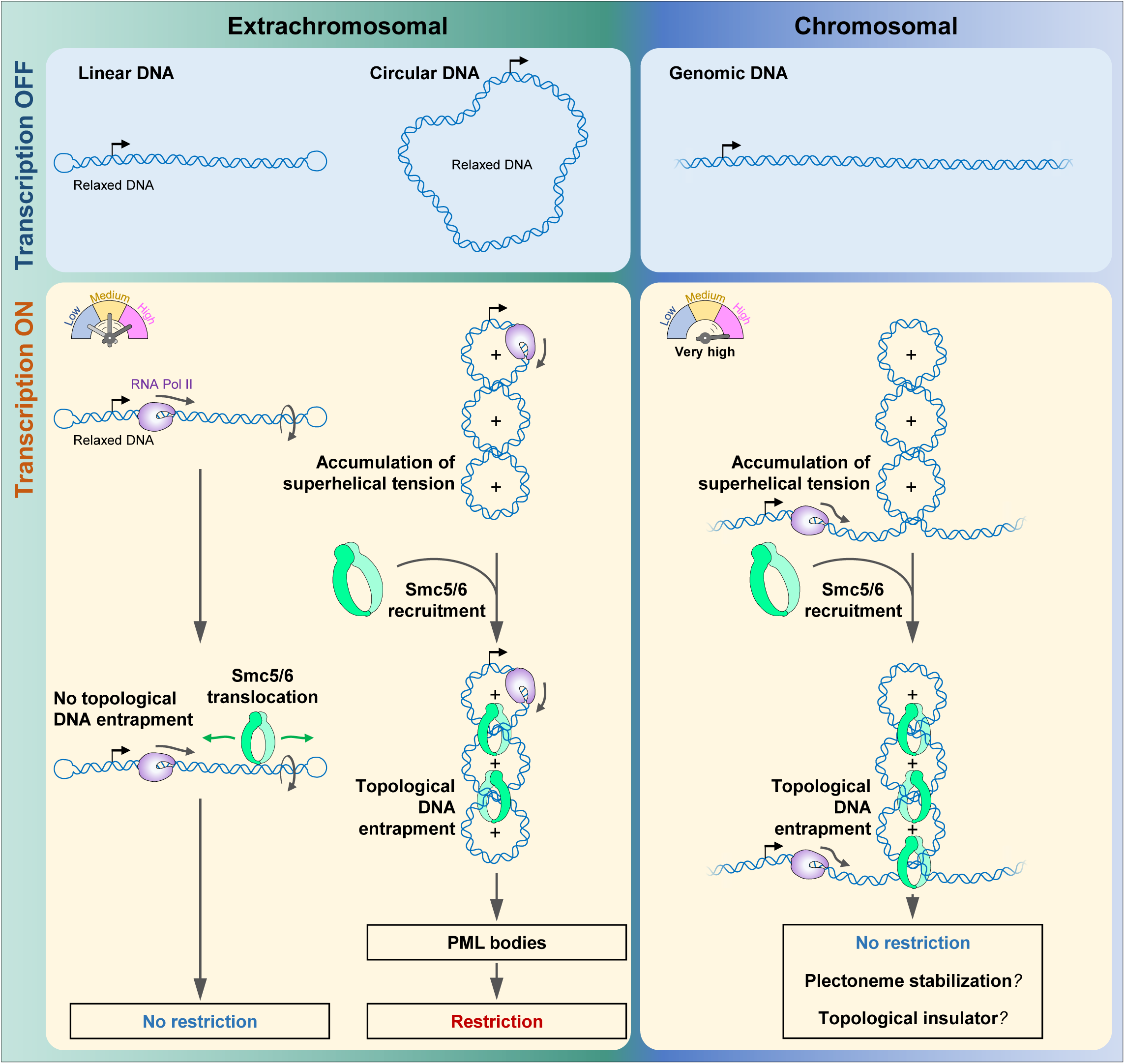
Speculative model. Upon transcription, the superhelical tension generated by the transcribing RNA polymerase II will be more significant on a covalently closed circular DNA molecule compared to a linear one, independently of the promoter strength. This will lead to the recruitment of Smc5/6 and to the topological DNA entrapment of the circular DNA while the complex will translocate along the linear DNA. On the chromosomes, an equivalent amount of superhelical tension can only be reached under conditions of a high transcriptional output, which also results in Smc5/6 recruitment. Based on previously reported data^30,37^, we hypothesize that, on chromosomes upon DNA entrapment, Smc5/6 stabilizes the plectonemic DNA structures formed. By preventing supercoil spreading, Smc5/6 would therefore act as a topological insulator and/or facilitate the resolution of such DNA structures. The same DNA entrapment on extrachromosomal circular DNA leads to ecDNA recruitment to PML bodies and restriction^23^.

The above results provide new insights into a long-standing controversy in the HBV field regarding the kinetics of HBx expression. How is HBx expressed from the viral covalently closed circular DNA (cccDNA) if Smc5/6 represses its transcription? Based on our data, we could envision a model in which the DNA supercoils that arise during the first round of transcription that enable HBx production, recruit Smc5/6 to silence the cccDNA which is then subsequently degraded by HBx to sustain effective viral transcription.

Our observations that Smc5/6 binding requirements are the same for both chromosomal and extrachromosomal DNA confirm that the restriction activity of the complex relies on subsequent steps. Further work is needed to decipher the detailed molecular events underlying the specificity of Smc5/6-mediated transcriptional repression towards extrachromosomal DNA.

## LIMITATIONS OF THE STUDY

Our study revealing the *in vivo* DNA topology-dependent association of Smc5/6 is inferred based on indirect evidence. This is due to the lack of efficient *in vivo* techniques permitting the high-resolution mapping of DNA supercoils at the individual gene level in mammalian cells. Moreover, as DNA supercoils are a combination of twist and writhe, we cannot exclude that Smc5/6 actually recognizes juxtaposed DNA helices, as previously suggested^30^. Therefore, future research should be aimed at developing new tools to monitor DNA supercoiling *in vivo*. Understanding of the molecular events leading to the specific Smc5/6-mediated transcriptional silencing of extrachromosomal DNA requires further characterization. Because ChIP-seq experiments do not provide temporal information regarding the dynamics of Smc5/6 recruitment and stabilization onto DNA, the use of single-molecule tracking combined with super-resolution microscopy would provide useful information to elucidate the Smc5/6-DNA interaction behavior.

## ACKNOWLEDGMENTS

We thank Patrick Meraldi (University of Geneva) and Howard Y. Chang (Stanford University) for sharing the hTERT-RPE and the COLO320DM cell lines. We thank Grzegorz Kudla (University of Edinburgh) for sharing T7 RNA polymerase plasmids and Mathieu Brochet (University of Geneva) for sharing the pJazz®-OK vector. We thank the *iGE3 Genomics Platform* of the University of Geneva for ChIP-Seq data (https://ige3.genomics.unige.ch) and the *Flow Cytometry facility* of the University of Geneva. We thank Fabien Abdul, Sari Kassem and the members of the Viollier laboratory (University of Geneva) for critical discussions. We are most grateful to Joseph Curran and Dominique Belin for critical reading of the manuscript. This work was supported by grants from the Swiss National Science Foundation 310030-149626 and 310030-175781 to M.S.

## AUTHOR CONTRIBUTIONS

Conceptualization of the project A.D. and M.S. A.D. designed and carried out all the experiments. A.D. G.P. and J.P. analyzed and interpreted the ChIP-Seq data. C.C. acquired and analyzed all the microscopy data. B.B. performed the western blots shown in Figures 3C, 3D, 4F and S4A. M.S. acquired the funding. A.D. and M.S. analyzed the data. A.D. wrote the manuscript with input from all authors.

## EXPERIMENTAL MODEL AND SUBJECT DETAILS

### Cell culture

The hTERT-RPE1 (non-transformed immortalized human retina pigment epithelial cells) (ATCC; CRL-4000) (kind gift from P. Meraldi), human embryonic kidney cell line HEK 293T/17 (ATCC; CRL-11268), human adenocarcinoma epithelial cells HeLa (ATCC; CCL-2) and the human colon cancer carcinoma COLO320DM (kind gift from H. Y. Chang) cells and their derivatives, were grown at 37°C under 5% CO2 in high glucose DMEM (Gibco; 41966029) supplemented with 10% FBS (Gibco; 10270-106), 1% penicillin/streptomycin (Sigma; P0781-100), 2 mM l-glutamine (25030024), 1 mM sodium pyruvate (11360070), and 1% MEM non-essential amino acids solution (11140050) (all from ThermoFischer).

To generate the hTERT-RPE1 cell line over-expressing the HA-tagged version of Smc6, hTERT-RPE1 cells (ATCC; CRL-4000) were transduced with pCDH-CMV-3xHA-Smc6-EF1α-RFP lentiviral vector. One month post transduction, positive RFP cells were FACS sorted and used for further experiments.

To obtain the hTERT-RPE1 clones with a chromosomally integrated excisable [GLuc^circle^] construct, hTERT-RPE1 cells (ATCC; CRL-4000) were transduced with a lentiviral vector (System Biosciences #CD511B) encoding the excisable reporter construct (see details below). Two weeks post transduction, single-cell clones were isolated by FACS in 96-wells plate and expanded for two more weeks before screening. GLuc and GFP expression, before and after Cre recombinase expression, was used as screening criteria for each clone.

### Transfections and treatments

siRNAs transfections were performed following the manufacturer’s instructions for 72 hours with 20 nM siRNAs using either siNTC or siTop1A, siTop2A and siTop2B (Table 1) (Horizon Discovery Ltd). Opti-MEM (ThermoFisher; 31985070) and Lipofectamine™ RNAiMAX (ThermoFisher; 13778150) were used.

**Table 1.**
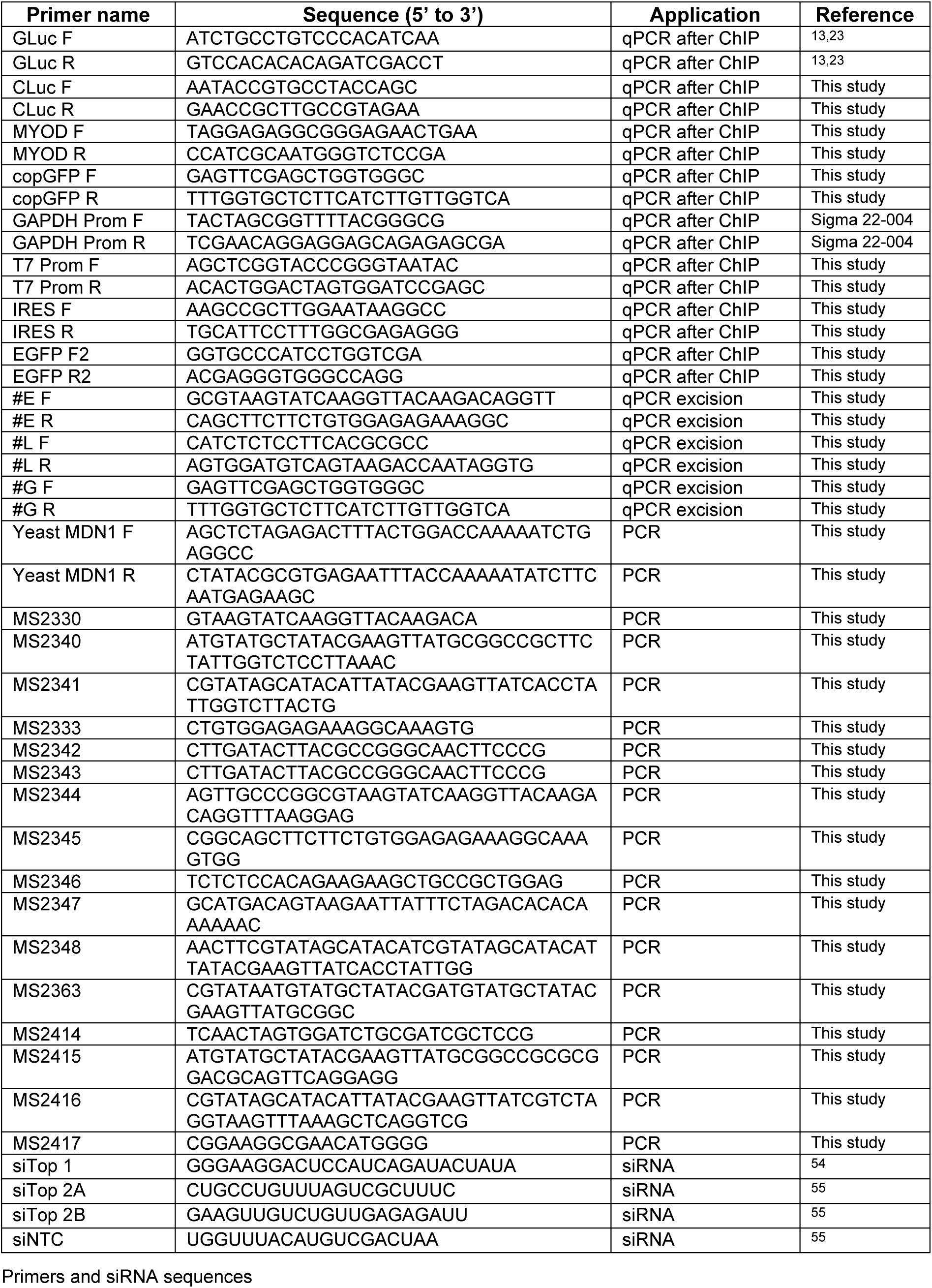
Primers and siRNA sequences

For plasmid transfection, cells were transduced with the appropriate lentiviruses 24 h prior transfection and seeded at a density of 6×10^6^ cells in a 10-cm dish. Cells were then reverse-transfected with 3 µg of reporter plasmid using Lipofectamine™ 2000 (ThermoFisher; 11668019) following the manufacturer’s instructions. Analysis was performed 48 h later.

Actinomycin D (Enzo Life Sciences; BML-GR300-0005) and triptolide (Sigma; T3652) were used as indicated on the figures.

## METHODS DETAILS

### Plasmids and Constructs

The lentiviral vector pWPT (Addgene #12255) was used to expressed either Mock (empty), GFP-HBx or HBx alone as described previously^13^. pCDH-CMV-MCS-EF1-Puro (System Biosciences #CD510B) was used to express HBx in case of experiments done in presence of Cre recombinase. The Cre recombinase was expressed from the pLOX-CW-CRE (Addgene #12238). pCDH-CMV-MCS-EF1α-RFP lentiviral vector (System Biosciences #CD512B) was used to clone the human codon-optimized, chemically synthesized (System Biosciences) Smc6 gene sequence under the control of the CMV promoter which was further 3xHA-tagged. The secreted Gaussia luciferase (GLuc) and the secreted Cypridina luciferase (CLuc) used as reporter were either expressed from pCDH-CMV-GLuc-EF1α-copGFP or pCDH-CMV-CLuc-EF1α-RFP, respectively, when delivered as extrachromosomal DNA using an integrase-defective (D116A) lentiviral vector (System Biosciences #CD511B).

The excisable reporter construct was generated through several steps. A first segment containing the split Gaussia Luciferase gene sequence (GLuc-Nter aa 1 to 49 and Gluc-Cter aa 50 to 186), obtained from pCMV-GLuc(M60I) and separated by a chimeric intron (splicing donor and acceptor sequences both from pCI-neo, Promega; E1731) with an embedded inverted *LoxP* site, was amplified using several rounds of overlapping PCR with primers pairs (MS2330-MS2340), (MS2341-MS2333), (MS2342-MS2343), (MS2344-MS2345), (MS2346-MS2347), (MS2342-MS2347) and cloned into the BamHI and XbaI sites of pCMV-empty to generate pCMV-GLuc(Nter)-donnor-LoxP-acceptor-GLuc(Cter).

A second fragment containing the EF1α-*LoxP*-copGFP sequence was amplified using several rounds of overlapping PCR from pCDH-CMV-GLuc-EF1α-copGFP with primers pairs (MS2414-MS2415), (MS2416-MS2417), (MS2414-MS2463) (MS2348-MS2417), and (MS2414-MS2417) and cloned into the SpeI and PstI sites of pCDH-CMV-GLuc-EF1α-copGFP to generate pCDH-EF1α-*LoxP*-copGFP.

A unique NotI restriction site was added in the *LoxP* site sequence to facilitate further cloning steps.

The *LoxP*-copGFP form pCDH-EF1α-*LoxP*-copGFP was cloned in the NotI and PstI sites from pCMV-GLuc(Nter)-donnor-*LoxP*-acceptor-GLuc(Cter) to generate pCMV-GLuc(Nter)-donnor-*LoxP*-copGFP.

In parallel a 3 kb DNA sequence from the budding yeast *MDN1* gene, later called DNA stuffer, was PCR amplified and cloned into the XbaI and MluI sites of pLVX-CMV-GLuc(M60I) to generate the pLVX-CMV-GLuc-DNAstuffer.

The *LoxP*-acceptor-GLuc(Cter) from pCMV-GLuc(Nter)-donnor-LoxP-acceptor-GLuc(Cter) was cloned in the NotI and XbaI sites of pLVX-CMV-GLuc-DNAstuffer. The GLuc(Nter)-donnor-*LoxP*-copGFP generated from pCMV-GLuc(Nter)-donnor-*LoxP*-copGFP was further cloned in the MluI and PstI sites from this newly created construct to generate pLVX-CMV-GLuc-*LoxP*-acceptor-GLuc(Cter)-DNA stuffer-CMV-GLuc(Nter)-donnor-*LoxP*-copGFP.

To generate the final lentiviral construct pCDH-EF1α-*LoxP*-acceptor-GLuc(Cter)-DNAstuffer-CMV-GLuc(Nter)-donnor-*LoxP*-copGFP, the *LoxP*-acceptor-GLuc(Cter)-DNAstuffer-CMV-GLuc(Nter)-donnor-*LoxP*-copGFP obtained after partial digestion of pLVX-CMV-GLuc-*LoxP*-acceptor-GLuc(Cter)-DNA stuffer-CMV-GLuc(Nter)-donnor-*LoxP*-copGFP was cloned in the NotI and PstI sites of pCDH-EF1α-*LoxP*-copGFP.

The linear pJazz mammalian expression vector was obtained by cloning the CMV-GLuc-EF1α-copGFP cassette from pCDH-CMV-GLuc-EF1α-copGFP into the SpeI and NotI sites of pJazz®-OK (kind gift from M. Brochet). Its circular counterpart was obtained after digestion with BssHII and self-ligation.

To generate the NLS-T7-polymerase lentiviral vector, the DNA fragment corresponding to the NLS-T7-polymerase sequence was PCR amplified using the primer pairs (5’-CGAACCCTTGGATCCGCCACCATG-3’ and 5’-GCCGCGGCCGCACCGGTAGGGATCG-3’) from the pcDNA3.4-T7pol plasmid (kind gift from G. Kudla)^56^ and cloned into the BamHI and NotI sites of the pCDH-CMV-MCS-EF1α-RFP lentiviral vector (System Biosciences #CD512B). The T7 promoter-IRES-GFP reporter lentiviral construct was generated by PCR amplification of the corresponding DNA fragment from the pUC19-T7-pro-IRES-EGFP plasmid (kind gift from G. Kudla)^56^ using the primer pairs (5’-CGGATCGATTGTAAAACGACGGCCAGTGAATTC-3’ and 5’-CGGTCGACTTAAAGACAGGCCTTACTGGCTGAATAGA-3’) and cloned into the ClaI and SalI sites of the pCDH-CMV-GLuc-EF1α-copGFP lentiviral vector (System Biosciences #CD511B) from which the CMV promoter was previously removed.

To generate the c-Myc expression vector, the c-Myc sequence from the pCDH-Puro-cMyc plasmid (Addgene #46970) was cloned into the XbaI and NotI sites of the pCDH-CMV-MCS-EF1α-RFP lentiviral vector (System Biosciences #CD512B).

The Myc-tagged DNA topoisomerase IB from vaccinia virus fused to a NLS (Myc-NLS-vTopIB) sequence was chemically synthesized and cloned in the sites XhoI and XbaI of the pLVX-CMV-MCS-PGK-Puro lentiviral vector (BioCat GmbH).

All the primers used are listed in Table 1. Phusion High-Fidelity DNA polymerase (Thermo Scientific; F530L) was used for all the PCR reactions. All the restriction enzymes and T4 DNA ligase were from NEB.

### Lentivirus Production and Titration

Briefly, 4.5×10^6^ HEK 293T/17 cells were plated in a 10-cm dish and transiently transfected 16 h later using the calcium phosphate method with 15 μg lentiviral vector plasmid, 10 μg of packaging plasmid psPAX2 (Addgene plasmid #12260), and 5 μg of envelope plasmid pMD2G (Addgene plasmid #12259) to produce VSV-G pseudotyped recombinant lentiviruses. To produce integrase-defective lentiviruses, the plasmid p8.9NdSB (king gift from J. Luban) which encoded a catalytically inactive integrase point mutant (D116A) was used in replacement of the psPAX2. 8 h post-transfection the culture medium was changed. Two days later, supernatants containing the viral particles were collected and filtered through PVDF 0.45 μm filters (Merck-Millipore; SLHV033RS) before storage at −80°C. The titre of lentiviruses expressing GFP or RFP was estimated, by FACS analysis of GFP-positive or RFP-positive cells after infecting HeLa cells for 4 days with serially diluted viral supernatants.

### Luciferase Reporter Gene Assay

Luciferase activities were measured 2-3 days after reporter transfection or transduction. Briefly, 5 µL of cell culture supernatants were mixed with 50 µL of Luciferase assay buffer (100 mM NaCl, 35 mM EDTA, 0.1% Tween^®^ 20, 300 mM sodium ascorbate in 1X phosphate-buffered saline) containing as substrate either: 4 μM coelenterazine (Biosynth AG; C-7001) in case of GLuc or 1 μM vargulin (NanoLight; 305) for CLuc. Luminescence was measured in triplicates in a 96 microplate luminometer (Glomax; Promega).

### Western Blotting

Cell lysis was performed in 2% sodium dodecyl sulfate (SDS) (Sigma; 71729) in water, and the lysates were denatured at 95°C for 5 minutes. Protein concentrations were estimated using the Pierce^TM^ BCA Protein Assay kit (Thermo Scientific; 23225). Equal amounts of proteins (30-50 μg) were separated on 8-18% SDS-PAGE gradient gels and electroblotted onto nitrocellulose membranes (Amersham; 10600003). The membranes were blocked in 5% (w/v) non-fat dry milk – PBST 0.1% [Phosphate-buffered saline (ThermoFisher; 14190094) supplemented with 0.1% Tween^®^ 20 (Sigma; P1379)] for 1h and incubated overnight with primary antibodies at 4°C in 5% milk – PBST 0.1%. The membranes were then washed thrice with PBST 0.1% for 10 min, and incubated with secondary antibodies for 1 h at RT. The membranes were finally washed thrice with PBST 0.1% for 10 min. Detection was performed with SuperSignal West Pico PLUS chemiluminiscent substrate (Thermo Scientific; 34580) according to the manufacturer’s protocol. Images were acquired with the ChemicDoc XRS luminescence imager from Bio-Rad. The primary and secondary antibodies used are listed in Table 2.

**Table 2.** Details of sources of antibodies used for immunoblotting and immunofluorescence in this study.

| **Antibody** | **Host animal** | **Distributor, cat #** | **Dilution for immunoblot** | **Concentration for immunostaining** |
| --- | --- | --- | --- | --- |
| Anti-Smc5 | Rabbit polyclonal | Bethyl A300236A | 1 :1000 | N/A |
| Anti-Nse4a | Rabbit polyclonal | Abcepta AP9909a | 1 :1000 | N/A |
| Anti-Top1 | Rabbit monoclonal | Abcam ab109374 | 1 :1000 | N/A |
| Anti-Top2a | Rabbit monoclonal | Abcam ab52934 | 1 :1000 | N/A |
| Anti-Top2b | Rabbit polyclonal | Abcam ab72334 | 1 :5000 | N/A |
| Anti-c-Myc | Mouse monoclonal | Covance MMS-150R | 1 :1000 | N/A |
| Anti-myc tag | Mouse monoclonal | DSHB 9E 10 | N/A | 1 :10 |
| Anti-T7 RNA pol | Rabbit polyclonal | Creative Diagnostics CABT-B8990 | N/A | 1 :100 |
| Anti-Hsp90 | Mouse monoclonal | SCBT sc-13119, F8 | 1 :5000 | N/A |
| Anti-GAPDH | Mouse monoclonal | SBCT sc-32233, 6C5 | 1 :3000 | N/A |
| Goat Anti-Mouse IgG (H + L)-HRP | Goat polyclonal | Bio-Rad 1706516 | 1 :10000 | N/A |
| Goat Anti-Rabbit IgG (H + L)-HRP | Goat polyclonal | Bio-Rad 1706515 | 1 :10000 | N/A |
| Donkey anti-Mouse IgG (H+L) Alexa Fluor 488 | Donkey polyclonal | ThermoFisher Scientific, A21202 | N/A | 1: 400 |
| Goat anti-Rabbit IgG (H+L) Alexa Fluor 488 | Goat polyclonal | ThermoFisher Scientific, A-11008 | N/A | 1: 400 |

### ChIP and Quantitative PCR

ChIP analysis was performed using chromatin extracted from about 5 × 10^6^ HA-tagged Smc6-expressing hTERT-RPE1 cells cultured in a 10-cm diameter dish. Cells were harvested with trypsin-EDTA and collected by low-speed centrifugation 500 x *g* for 5min. Cells were rinsed once with PBS, resuspended in DMEM and fixed with 1% formaldehyde (Sigma; 47608) for 10 min at RT before quenching with 330 mM glycine 5 min at RT and then 15 min on ice. After two further washes with ice-cold PBS, cells were resuspended and lysed for 10 min at 4°C in 1 mL Cell Lysis Buffer (20 mM Tris-HCl (pH 8.0), 85 mM KCl, 0.5% NP-40) supplemented with EDTA-free protease inhibitor cocktail (Roche; 4693132001). The nuclei were collected by centrifugation at 500 x *g* for 5 min at 4°C and washed once in the same buffer. Nuclei were resuspended in 500 μL Nuclei Lysis Buffer (10 mM Tris-HCl (pH 7.5), 1% NP-40, 0.5% sodium deoxycholate, 0.1% SDS, protease inhibitor cocktail) and incubated for 10 min at 4°C. Chromatin was fragmented by sonication 3 × 10 s at 60% duty cycle, with 30 sec on wet-ice between sonication cycles, using a microtip-equipped SLPe sonifier (Branson Ultrasonics ™ Sonifier™, Brookfield, USA). The sonicated samples were centrifuged at 16,000 x *g* for 10 min. The supernatants were collected and 50 µL (1/10) was set aside as input controls. The rest (450 µL) was diluted with 1500 μl ChIP Dilution Buffer (0.01% SDS, 1.2 mM EDTA, 1.1% Triton X-100, 16.7 mM Tris-HCl (pH 8), 167 mM NaCl) supplemented with protease inhibitors and mixed with 50 μL Dynabeads™ Protein G (Invitrogen; 10009D) coupled either to 3 μg anti-HA antibody (Invitrogen; MA5-27915) in case of HA-Smc6 analysis, 5 μg anti-RPB1 (8WG16) (Covance; MMS-126R) for RNA pol II, 2 μg anti-H3pan (Diagenode; C15410324) or 4 μg anti-H3.3 (Diagenode; C15210011). After overnight incubation at 4°C on a rotating wheel, the beads were washed twice with 1 mL Cell Lysis Buffer, twice with high-salt buffer (50 mM HEPES-KOH (pH 7.5), 500 mM NaCl, 1 mM EDTA, 1% Triton X-100, 0.1% sodium deoxycholate), once with LiCl buffer (10 mM Tris-HCl (pH 8.0), 1 mM EDTA, 250 mM LiCl, 1% NP-40, and 1% sodium deoxycholate) and once with TE buffer (10 mM Tris-HCl (pH 8.0), 1 mM EDTA). To elute immunoprecipitated chromatin fragments, beads were incubated for 10 min at 65°C in 400 μL freshly prepared elution buffer (1% SDS, 0.1 M NaHCO_3_). DNA crosslinks were reversed by overnight incubation at 65°C with 0.6M NaCl and 80 μg proteinase K (Eurobio Scientific; GEXPRK01-I5). Samples were extracted once with Phenol – chloroform – isoamyl alcohol (Sigma; 77617), once with chloroform (Reactolab SA; P02410E16), ethanol precipitated and then resuspended in water. The input DNA samples were treated identically. The recovered DNA were quantified by real-time PCR using the KAPA SYBR FAST qPCR Kit Master Mix (2X) Universal (Roche; SFUKB) and the Bio-Rad CFX96 Real-time PCR System. The values shown in the figures are the ratios between the ChIP signals and the respective input DNA signals. The oligonucleotide primers used are listed in Table 1.

### ChIP-seq

ChIP-Seq DNA was purified as it was described in ChIP and Quantitative PCR paragraph. ChIP-enriched DNA were used to prepare libraries and processed with the Illumina TruSeq ChIP kit according to manufacturer specifications. Library molarity and quality were assessed with the Qubit (Thermofisher Scientific) and Tapestation (Agilent Technologies - DNA High sensitivity chip). Libraries were sequenced on a HiSeq 4000 and a NovaSeq 6000 Illumina sequencers for SR100 reads. Bioinformatics processing: sequenced reads were aligned to the human genome (build GRCh38.p13 downloaded from ENCODE) augmented with GLuc_episomal DNA sequence. We used the “mem” command of the software BWA (v0.7.17) for the alignment [BWA]^57^, and convert the output into sorted BAM with SAMTOOLS v1.10. ChIP-seq peak calling was performed with MACS2 (v2.2.7.1)^58^ command “callpeak” and parameters “--format BAM --gsize hs --SPMR --keep-dup 1 --qvalue 0.05 --nomodel --extsize 1000” [MACS]. MACS2 is used on one hand to detect Smc5/6 peaks comparing DMSO to No-HA reference condition; and on the other hand to detect RNA pol II peaks comparing DMSO to Input condition.

Coverage computations and visualizations were done using scripts in R programming language making use of Bioconductor package. The code consisted in loading Smc5/6 peaks detection of MACS2 and retain those with a fold enrichment above 2.5. Smc5/6 peaks were annotated with genes (taken from GENECODE v41 GTF) located within a 10kb region of the peaks and curated manually. GenomicRanges R package and findOverlaps() method is used in this process. Then, Smc5/6 peaks and RNA pol II peaks were merged and reduced (R method GenomicRanges ::reduce()) to determine genomic regions where they overlap (findOverlaps()).

Transcriptomic expression for RPE1 cell line was retrieved from GSE89413^47^, and we consider FPKM normalized values of each gene to sort them by expression. We matched genes with Smc5/6 peaks by names.

To produce the heatmaps, we resized the ChIP-Seq mapped reads to 1kb length, and compute at each genomic position the amount of overlap between forward-stranded and reverse-stranded reads (=2.min(fwd,rev)) normalized as read-per-million. A similar scaling factor is then applied to each gene of all condition so gene profiles can be compared from one gene to the other.

### DNA extraction and qPCR

Cells were lysed in DNA lysis buffer (100 mM Tris-HCl (pH 7.5), 10 mM EDTA, 1% SDS) and briefly sonicated. Samples were extracted once with Phenol – chloroform – isoamyl alcohol (Sigma; 77617), once with chloroform (Reactolab SA; P02410E16), ethanol precipitated and then resuspended in water. The recovered DNA were quantified by real-time PCR using the KAPA SYBR FAST qPCR Kit Master Mix (2X) Universal (Roche; SFUKB) and the Bio-Rad CFX96 Real-time PCR System. The oligonucleotide primers used are listed in Table 1.

### EU labelling

Transcription inhibition was monitored through EU incorporation into nascent RNAs using Click-iT RNA Alexa Fluor 488 Imaging Kit (Invitrogen; C10329) according to the manufacturer’s instructions. Briefly, after treatment as indicated in figure legends, hTERT-RPE1 cells seeded onto glass coverslips were incubated with 1 mM EU for 1h. The cells were fixed with 3.7% formaldehyde in PBS for 15min at RT and then permeabilized with 0.5% Triton X-100 for 15 min at RT. Incorporated EU was labelled by Click-iT reaction according to the manufacturer’s instructions for 30 min at RT. The cells were washed with PBS before mounting the coverslips with VECTASHIELD containing DAPI (Vector Laboratories, H-1200). Immunofluorescence images were acquired on an Olympus DeltaVision wide-field microscope (GE Healthcare) equipped with a DAPI/FITC/TRITC/Cy5 filter set (Chroma Technology Corp.) and a Coolsnap HQ2 CCD camera (Roper Scientific) running Softworx 6.5.2 (GE Healthcare). 3D images were deconvolved using Softworx 6.5.2 (GE Healthcare). Fixed cells were imaged with a 40x NA 1.35 oil objective in 0.2 µm Z-stacks.

### Live-cell imaging and Immunofluorescence

For live-cell imaging of the hTERT-RPE1 cells containing the genomic excisable construct [GLuc^circle^], cells were seeded in a glass bottom height-well µ-Slide Ibidi chamber (Ibidi; 80826) and co-transduced with lentiviruses containing either no gene insert (Mock) or HBx, plus or minus the Cre recombinase for 48 h. 4 h prior live-cell imaging, the culture medium was replaced by Leibovitz L-15 medium (ThermoFisher; 21083) supplemented with 10% FBS and 1% penicillin/streptomycin and containing 25 nM SiR-DNA (Spirochrome; SC007). Cells were acquired on a Nikon Eclipse Ti-E wide-field microscope (Nikon) equipped with a DAPI/FITC/Rhod/CY5 filter set (Chroma Technology Corp.), an Orca Flash 4.0 CMOS camera (Hamamatsu), and an environmental chamber using NIS software (Nikon). Z-stacks were imaged with z-slices separated by 2.5 µm, with 50 ms exposure per z-slice in the GFP and Cy5 channels using a 40x NA 1.3 oil objective and 2 × 2 binning.

For fixed-cell imaging, hTERT-RPE1 cells over-expressing HA-tagged version of Smc6 and either T7 RNA pol or Myc-NLS-vTop1B were grown onto acid-etched glass coverslips. Cells were fixed with 3.7% formaldehyde for 15min before permeabilization with 0.5% Triton X-100 for 15min followed by blocking for 1 h in PBS supplemented with 3% BSA. The final dilution of primary antibodies were 1:100 for anti-T7 RNA Polymerase (Creative Diagnostics; CABT-B8990) and 1:10 for anti Myc-Tag (DSHB; 9E 10). Secondary antibody conjugated with Alexa Fluor 488 either goat anti-rabbit IgG (Invitrogen; A-11008) or Alexa Fluor 488 donkey anti-mouse IgG (Invitrogen; A21202) were used at 1:400. All antibodies were diluted in PBS supplemented with 3% BSA. Primary and secondary antibodies were applied for overnight and 60 min, respectively. Immunolabelled cells were washed with PBS before mounting the coverslips with VECTASHIELD containing DAPI (Vector Laboratories; H-1200). Immunofluorescence images were acquired on an Olympus DeltaVision wide-field microscope (GE Healthcare) equipped with a DAPI/FITC/TRITC/Cy5 filter set (Chroma Technology Corp.) and a Coolsnap HQ2 CCD camera (Roper Scientific) running Softworx 6.5.2 (GE Healthcare). 3D images were deconvolved using Softworx 6.5.2 (GE Healthcare). Fixed cells were imaged with a 40x NA 1.35 oil objective in 0.2 µm Z-stacks. Alternatively, fixed cells expressing Myc-NLS-vTop1B were visualised with a Plan Apo 40x (NA 1.3 Oil DICIII) objective in 0.2 µm Z-stacks on a spinning disk microscope (Nipkow Disk) Zeiss Cell Observer.Z1 equipped with a HXP120 fluorescence wide-field visualisation lamp and with a CSU X1 automatic Yokogawa spinning disk head. 512 × 512 pixel images were acquired with an Evolve EM512 camera and Visiview 4.00.10.

## Statistical analysis

The statistical analysis was performed using GraphPad Prism 8 (GraphPad). The statistical tests employed are described in the figure legends.

## Code and data availability

All deep sequencing data reported in this paper (ChIP-seq) were deposited in the GEO repository with accession number (GSEXXXXXX) and are publicly available as of the date of publication. This paper does not report original code since it is based on implementation of publicly available software. Published software and code used in this study are listed in the Key resource table. Any additional information required to reanalyse the data reported in this paper is available from the lead contact upon reasonable request.

