## Supplemental Figures for "Human Smc5/6 recognises transcription-generated positive DNA supercoils"

### SUPPLEMENTAL FIGURE LEGENDS

**Figure S1. Both non-replicated and replicated extrachromosomal DNA are constrained by Smc5/6**

(A) Schematic diagram illustrating the position of the primer pairs used to measure Cre/*loxP* recombination excision efficiency. Primers used for PCR amplification labeled in this figure are described in Table 1.

(B) qPCR Quantification of Cre/*loxP* -mediated recombination.

(C) Luciferase reporter assay for two other clonal cell lines of hTERT-RPE1 containing the genomic excisable [GLuc$^{circle}$] construct (clones 2 and clone 3). Same experimental setup and statistical analysis as in Fig 1C. Data are means ± SEM of 3 independent experiments. Statistical analysis was performed using one-way ANOVA with Tukey's multiple comparisons.

(D) Same results as in Fig 1D. Histone binding at the indicated loci was monitored by ChIP using anti-H3 or anti-H3.3 antibodies. Values are expressed as a percentage of input DNA recovered. Data are means ± SEM of 3 independent experiments.

**Figure S2. Circular DNA templates are the preferential target of Smc5/6**

(A) $C_t$ value for the extrachromosomal GLuc gene obtained with primers as listed in table Table 1 from DNA extracted from hTERT-RPE1 cells treated as in Fig 2B. Data are means ± SEM of 3 independent experiments. Statistical analysis was performed using the Student's t-test.

(B) Luciferase reporter assay, same sample as in Fig 2B. Luciferase activity was presented as relative luminescence units (RLU). Data are means ± SEM of 3 independent experiments.

**Figure S3. Time course imaging of transcription inhibition using 5-ethynyl uridine incorporation and visualization by click chemistry.**

Validation of the transcription inhibition in hTERT RPE-1 cells treated with (A) 10 µg/ml Actinomycin D (ACTD) or (B) 10 µM Triptolide (TPT), for 0-2-5-24h. Nascent RNA synthesis was monitored by the incorporation of an analog of uracil, 5-ethynyl uridine (5-EU), and visualized with Alexa Fluor 488 labeling by click chemistry. Nuclei were stained with DAPI. Scale bar, 50 µm.

**Figure S4. Topoisomerase levels in COLO320DM**

(A) Western blot showing topoisomerase 1 (Top1), topoisomerase 2A (Top2A) and topoisomerase 2B (Top2B) protein levels in protein extracts from COLO320DM cells transfected with either a non-targeting control siRNA (siNTC) or with siRNAs against topoisomerase 1 (siTop1) and topoisomerases 2A and

2B (siTop2). The Western blot analysis on COLO320DM cells was used to validate the siRNA KD and the Top2A antibody.

**Figure S5. Correlation between Smc5/6 and RNA pol II binding sites**

(A-B) Fold enrichment of reads mapping on (A) extrachromosomal GLuc coding sequence (558 bp) or (B) Chromosomal RFP coding sequence (681 bp) from HA-Smc6 ChIP-seq (blue bars) or RNA Pol II ChIP-seq (purple bars) samples.

(C) RNA-seq data from hTERT RPE-1 cells (GSE89413) showing the correlation between the highly transcribe genes and the presence of Smc5/6 peaks. Genes (x-axis) are ranked according to their expression level in FPKM (y-axis). Smc5/6 peaks are depicted in blue.

(D) Heatmaps of RNA pol II ChIP-seq peaks distribution in hTERT-RPE1 cells over-expressing a HA-tagged version of Smc6 treated with either DMSO (DMSO), the corresponding input (Input NT) or 10 μM Triptolide (TPT) for 24h before RPB1-ChIP. Cells were transduced with an integrase-defective lentiviral luciferase reporter construct (GLuc). Rows represent RNA pol II binding sites ±2 kb around the RNA pol II peak summit ranked by signal intensity (left panel). The color scale represents the ChIP-seq normalized read depth (RPM) row-scaled identically across the 3 samples, with mapped reads virtually resized to 1 kb-length and looking at each genomic position for the amount of overlap between forward- and reverse-stranded reads. On the right-hand panel, the blue dots represent RNA pol II peaks whose location match with a Smc5/6 peak (blue dots at the bottom are Smc5/6 peaks not matching any detected RNA pol II peak).

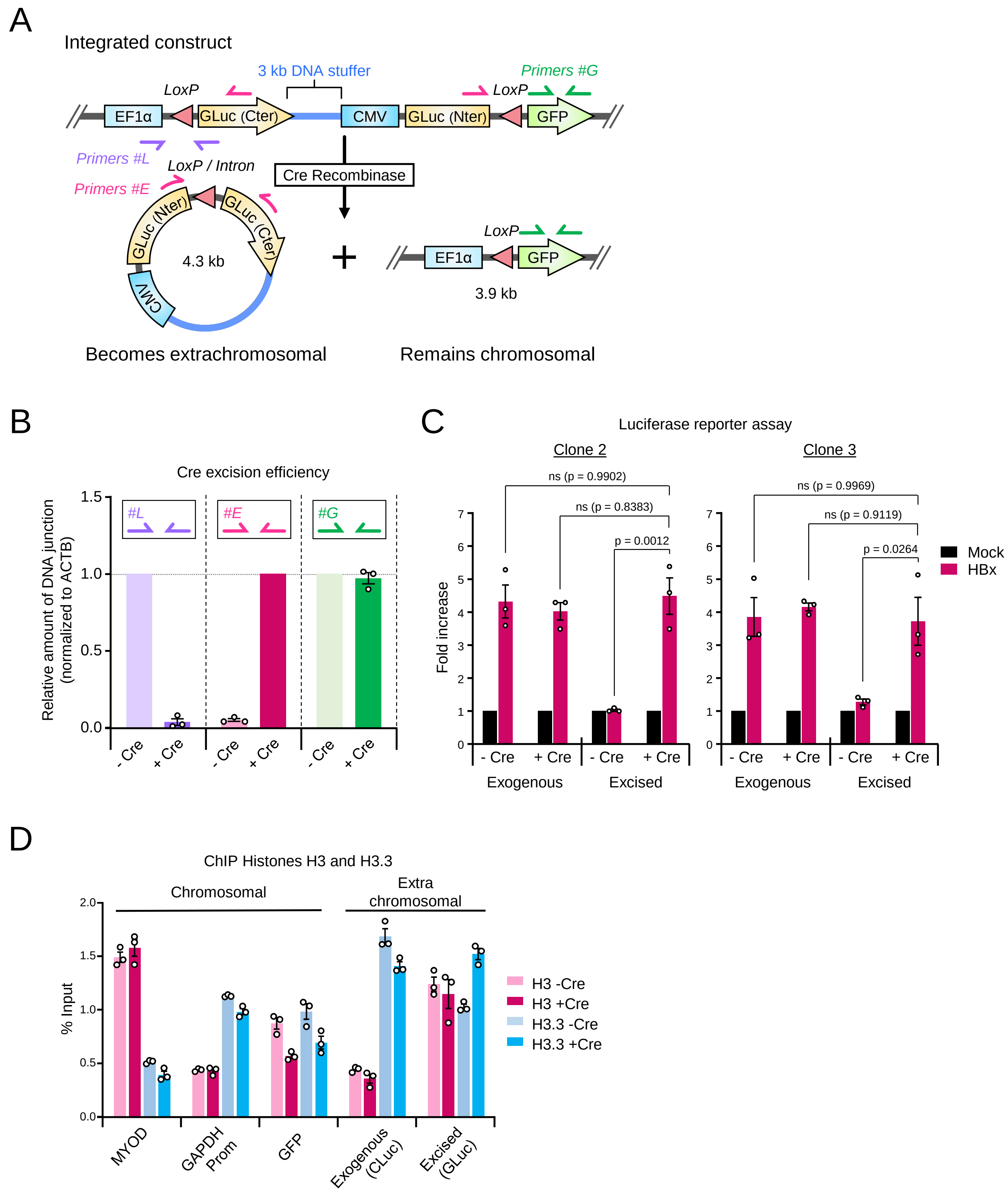


**Figure 1. Both non-replicated and replicated extrachromosomal DNA are constrained by Smc5/6**

Supplementary Figure 2 Diman et al.

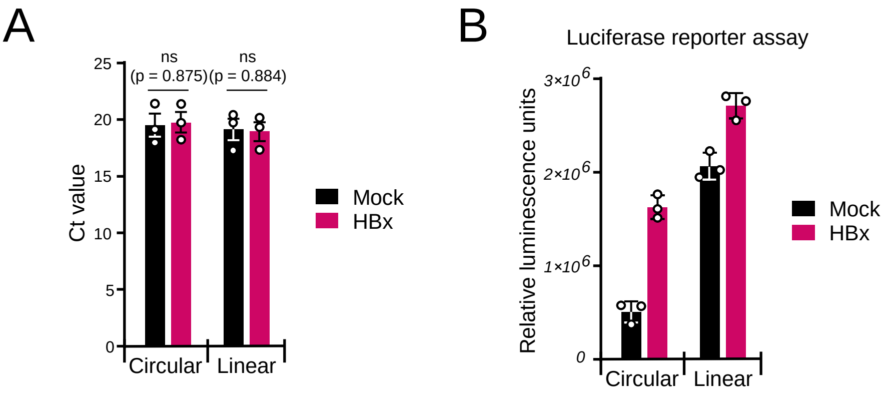


**Figure S2. Circular DNA templates are the preferential target of Smc5/6**

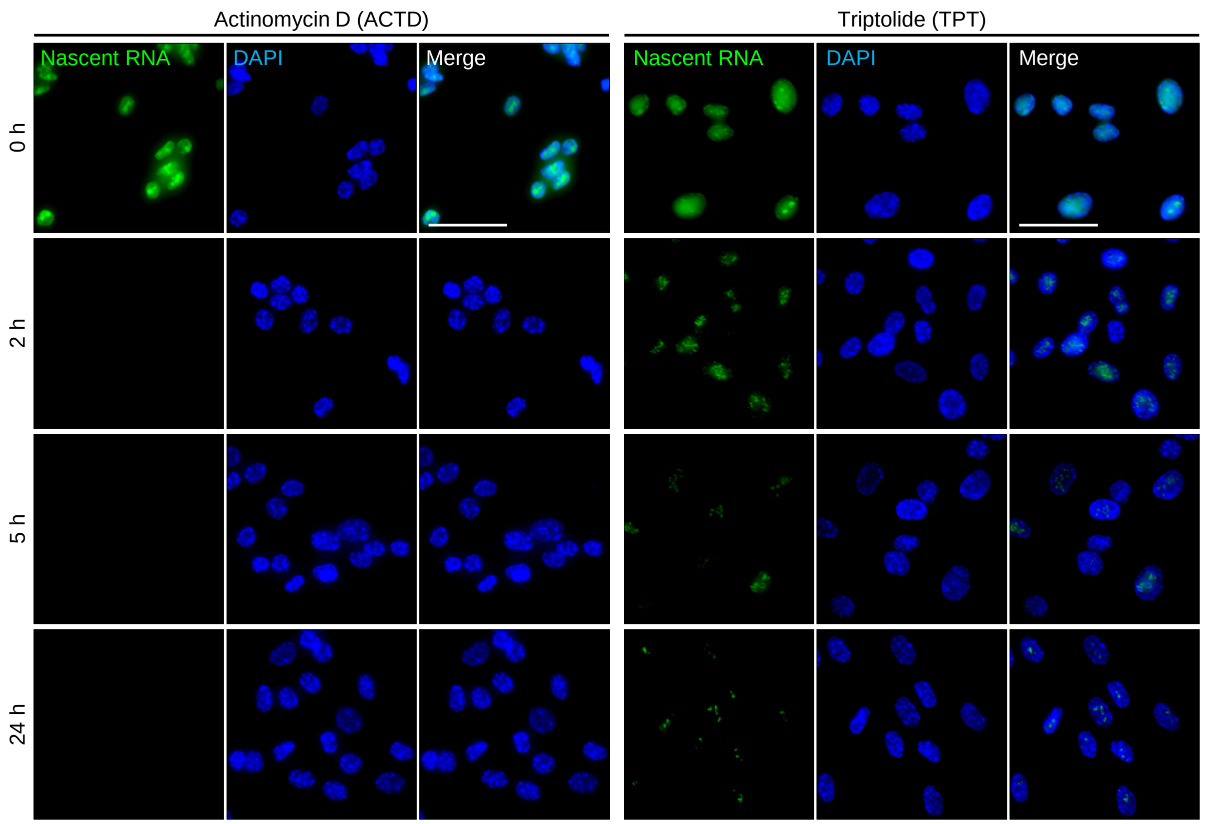


**Figure S3. Time course imaging of transcription inhibition using 5-ethynyl uridine incorporation and visualization by click chemistry.**

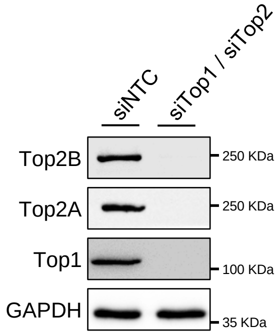


**Figure S4. Topoisomerase levels in COLO320DM**

Supplementary Figure 5 Diman et al.

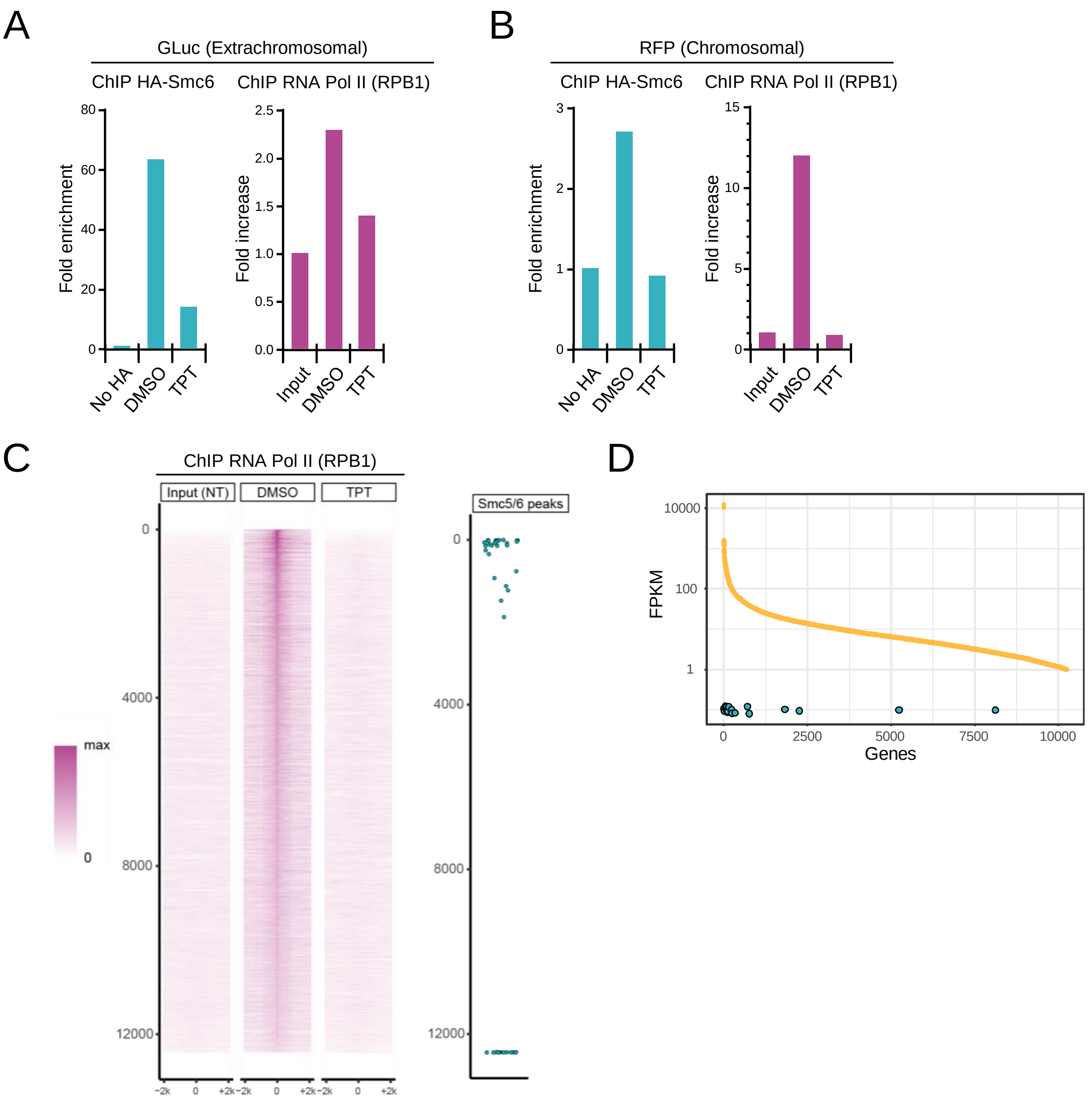


**Figure S5. Correlation between Smc5/6 and RNA Pol II binding sites**
